# Marginal specificity in protein interactions constrains evolution

**DOI:** 10.1101/2023.02.18.529082

**Authors:** Dia A. Ghose, Kaitlyn E. Przydzial, Emily M. Mahoney, Amy E. Keating, Michael T. Laub

## Abstract

The evolution of novel functions in biology relies heavily on gene duplication and divergence, creating large paralogous protein families. Selective pressure to avoid detrimental cross-talk often results in paralogs that exhibit exquisite specificity for their interaction partners. But how robust or sensitive is this specificity to mutation? Here, using deep mutational scanning, we demonstrate that a paralogous family of bacterial signaling proteins exhibits marginal specificity, such that many individual substitutions give rise to substantial cross-talk between normally insulated pathways. Our results indicate that sequence space is locally crowded despite overall sparseness, and we provide evidence that this crowding has constrained the evolution of bacterial signaling proteins. These findings underscore how evolution selects for ‘good enough’ rather than optimized phenotypes, leading to restrictions on the subsequent evolvability of paralogs.

**Significance Statement:** Large paralogous protein families are found throughout biology, the product of extensive gene duplication. To execute different functions inside cells, paralogs typically acquire different specificities, interacting with only desired, cognate partners and avoiding cross-talk with non-cognate proteins. But how robust is this interaction specificity to mutation? Can individual mutations lead to cross-talk or do paralogs diverge enough such that multiple mutations would be required, providing a mutational ‘buffer’ against cross-talk? To address these questions, we built mutant libraries that produce all possible single substitutions of a bacterial kinase and then screened for cross-talk to non-cognate proteins. Strikingly, we find that many single substitutions can produce cross-talk, meaning that these pathways typically exhibit only ‘marginal specificity’, and demonstrate that this restricts their evolvability.

## Introduction

The process of gene duplication and divergence fuels the evolution of proteins with new functions^1^. This fundamental mechanism has created large paralogous protein families within all clades of life^2,3^. However, the expansion of these protein families presents a challenge when members are required to bind distinct interaction partners^4–8^. Given their highly similar structures and sequences, how do the individual members of such families maintain different interaction specificities? And, do paralogs constrain each other’s evolution?

Answers to these questions lie in the nature of the sequence space relevant to such paralogous families. This sequence space is defined by the set of residues governing the interaction specificity of paralogs and their binding partners. In sequence space, each paralog must reside within a specific ‘niche’, defined here as the set of sequences capable of interacting with its binding partner(s). A given paralog may also have to avoid the niches of other proteins within this space to maintain interaction specificity. How much constraint is posed by other paralogs depends on the size, distribution, and extent of overlap of niches within sequence space.

Prior work demonstrated that the sequence space of some paralogous protein families is sparsely occupied, with ample room for new members, based on the observation that new, synthetic proteins could be readily discovered or introduced without cross-talk to existing systems^9–12^. However, the overall distribution of niches for extant paralogs in sequence space is not known, and there are two general possibilities. First, niches could be widely distributed throughout sequence space. Due to either selection pressure^13–18^ or the drift of sequences over evolutionary time, individual niches may have moved far apart in space, resulting in “robust specificity” in the sense that cross-talk between paralogs would require multiple substitutions (Fig. 1A, top). Alternatively, niches for extant paralogs could be clustered and partially overlapping (Fig. 1A, bottom), creating crowded local regions of sequence space despite overall sparsity. This could result in “marginal specificity” such that individual substitutions could, in principle, produce cross-talk. Such marginal specificity is akin to the well-documented marginal stability of proteins in which proteins are often just above a threshold stability level needed for folding^19–23^. This marginal stability arises because evolution does not select for additional stability once a protein can stably fold. A consequence, or reflection, of marginal stability is that individual substitutions can lead to unfolding. Marginal specificity could similarly arise if recently duplicated paralogs are under selective pressure only to separate in sequence space just enough to prevent unwanted cross-talk, with no pressure to further diverge and enhance the robustness of specificity.

**Fig. 1.**
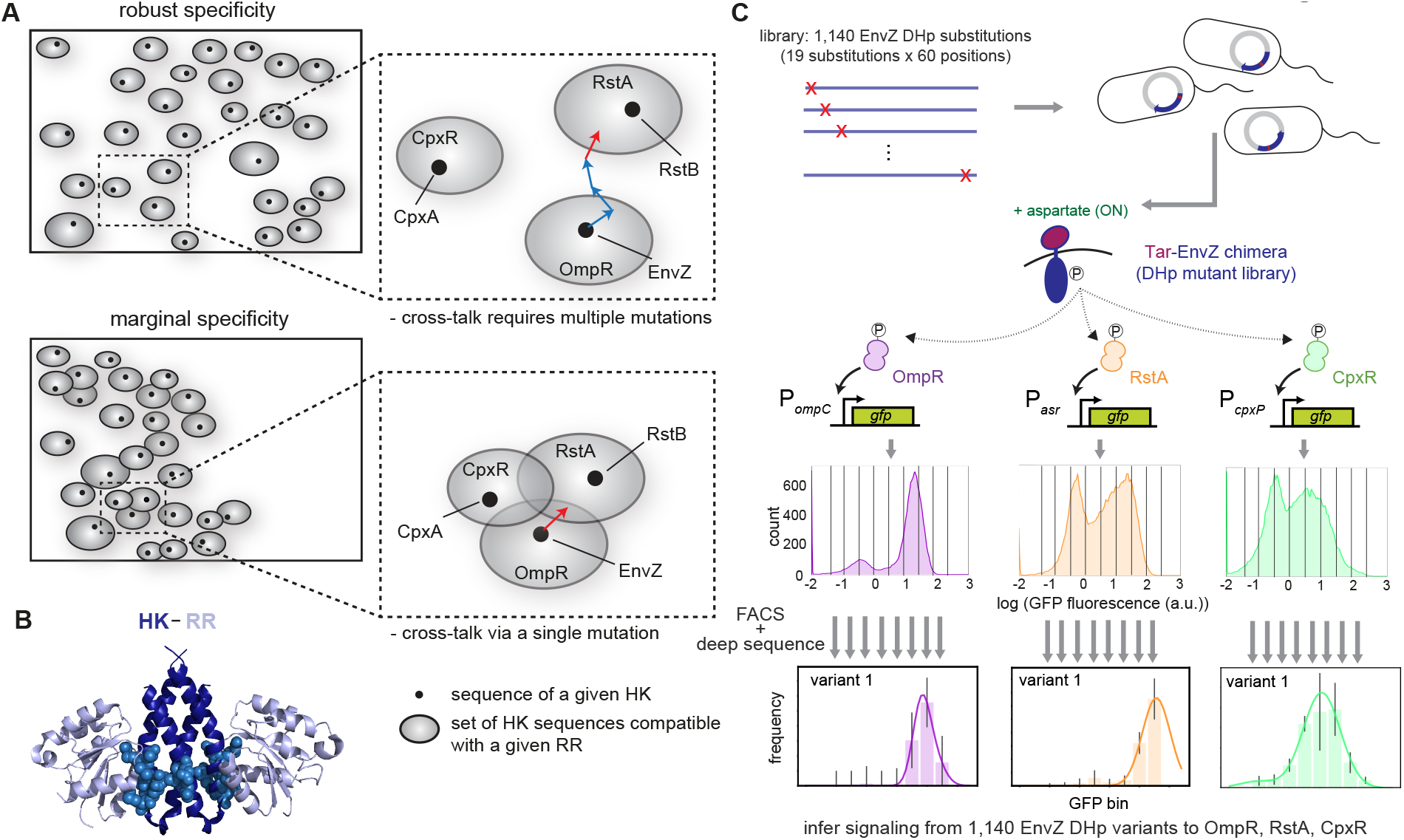
Assessing the density of local sequence space. (A) Sequence space diagram in which rectangles represent the space of all possible histidine kinase (HK) sequences, dots represent extant paralogs in a species (HKs in *E. coli*), and grey spheres represent the set of HK sequences that interact with a given response regulator (RR). HK paralogs must interact with their cognate RR, *i.e*., be within the niche of that RR, but avoid the niches of non-cognate RRs. Top: robust specificity model, where niches are generally well separated in sequence space. Bottom: marginal specificity model, where niches are separated just enough to avoid cross-talk, but often overlap such that local sequence space can be crowded. Insets show the arrangement of niches for OmpR, RstA and CpxR. In the robust specificity model, multiple mutations (depicted as arrows) to an HK such as EnvZ are required to introduce cross-talk; in the marginal specificity model, a single mutation may introduce crosstalk. (B) Model of an HK-RR complex. RR chains (light blue) from PDB ID 3DGE are positioned relative to the structure of EnvZ (5B1N, deep blue) using the DHp domains in each structure for alignment. HK positions that co-evolve with positions in the RR are shown as sky blue spheres. (C) A library of single-substitution EnvZ variants was transformed into three GFP reporter strains that read out activation of OmpR (P_*ompC*_), RstA (P_*asr*_), or CpxR (P_*cpxP*_). The resulting populations showed a distribution of GFP fluorescence and were sorted into 8 bins based on GFP level. Populations from each bin were deep sequenced, the frequency of variants in each bin calculated, and profiles fit to Gaussians to extract the peak fluorescence of each variant. Error bars in frequency profiles indicate standard deviation from 3 replicates.

To distinguish between these models of specificity, we investigated two-component signaling pathways, the most prevalent form of signal transduction system in bacteria, with most species encoding dozens of paralogous pathways^24^. The typical pathway consists of a histidine kinase (HK) that detects a signal, autophosphorylates, and then transfers a phosphoryl group to a co-operonic, cognate response regulator (RR). The phosphorylated RR elicits a cellular output, often by regulating gene expression^25^. The HK also dephosphorylates its cognate RR in the absence of signal^26^. There is typically very high structural and sequence similarity between paralogs in the domains responsible for HK-RR interactions, the dimerization and histidine phosphotransfer (DHp) domain of the HK and the receiver (REC) domain of the RR^27^. However, interactions between cognate HK-RR pairs are highly specific *in vivo* and *in vitro*, with little cross-talk between non-cognate partners^28^. HK-RR specificity is determined primarily by a subset of residues that strongly coevolve (Fig. 1B and S1A)^29,30^. These residues are found at the HK-RR interface and when collectively swapped from one system to another, they are often sufficient to rewire interaction specificity^29,20^. Introducing multiple substitutions at these key residues can produce cross-talk between non-cognate proteins that is severely detrimental to cellular fitness in certain conditions^4^. However, how likely individual substitutions are to produce cross-talk has not been systematically probed. Thus, whether two-component signaling pathways exhibit marginal or robust specificity is not yet known.

## Results

### A high-throughput method for assessing cross-talk between signaling pathways

To assess paralog specificity and determine how crowded sequence space is locally, we focused on the *E. coli* two-component signaling systems EnvZ-OmpR, RstB-RstA, and CpxA-CpxR (Fig. 1A). These three systems are widespread in β- and γ-proteobacteria, likely resulting from two ancient duplication and divergence events that occurred ~2 billion years ago in their common ancestor^31^ (Fig. S1B and S1C). To examine the occupancy of sequence space immediately surrounding EnvZ, we sought to measure the ability of variants harboring each possible single substitution in the DHp domain to activate the cognate regulator OmpR and the non-cognate regulators RstA and CpxR. To monitor activation, we generated strains in which a GFP reporter is expressed from a known OmpR-, RstA-, or CpxR-regulated promoter^32–34^ (Fig. 1C and S2A). Because the native signal for EnvZ is not known, we deleted *envZ* in each strain and introduced *taz*, which encodes a chimeric receptor containing the aspartate-sensing domain of the chemoreceptor Tar fused to the cytoplasmic, signaling domains of EnvZ^33,35,36^ (Fig. 1C). Taz drives robust (~14-fold), signal-dependent induction of our OmpR reporter, but not the RstA or CpxR reporters, reflecting the limited cross-talk between the three wild-type pathways (Fig. S2B and S2C). For simplicity, we refer to the wild-type Taz construct as EnvZ. To assess cross-phosphorylation of the non-cognate regulators RstA and CpxR, we deleted *rstB* and *cpxA* from each strain to prevent the phosphatase activity of these kinases from counteracting any phosphorylation of RstA or CpxR by EnvZ^33^ (Fig. S2A).

To focus our investigation on the local sequence space immediately surrounding EnvZ, we performed deep mutational scanning^37,38^. We constructed a library of all 1,140 single substitutions in the 60 amino-acid DHp domain of EnvZ, and then used a high-throughput screening approach, Sort-seq^9^, to measure interaction with OmpR, RstA, and CpxR (Fig. 1C). We transformed the library into each reporter strain, and then grew cells in the presence or absence of signal (aspartate) for 3 hrs before sorting cells into bins based on their fluorescence. The plasmids encoding *envZ* in cells from each bin in each condition were deep sequenced to determine the frequency of each variant, with frequency profiles fit to a Gaussian to extract the peak fluorescence values^9^ (Fig. 1C, S2D–S2J). To validate these values, 30 variants spanning the range of output fluorescence values for each regulator in both conditions were measured individually using flow cytometry and the median fluorescence compared to the values obtained from Sort-seq. For each reporter, there was a high correlation (R^2^ > 0.8) between the values obtained from Sort-seq and flow cytometry (Fig. 2A). We also purified a selection of 14 EnvZ variants and used ^32^P-based phosphotransfer assays to demonstrate that the activities towards the regulators seen *in vivo* were recapitulated *in vitro* (Fig. S3A).

**Fig. 2.**
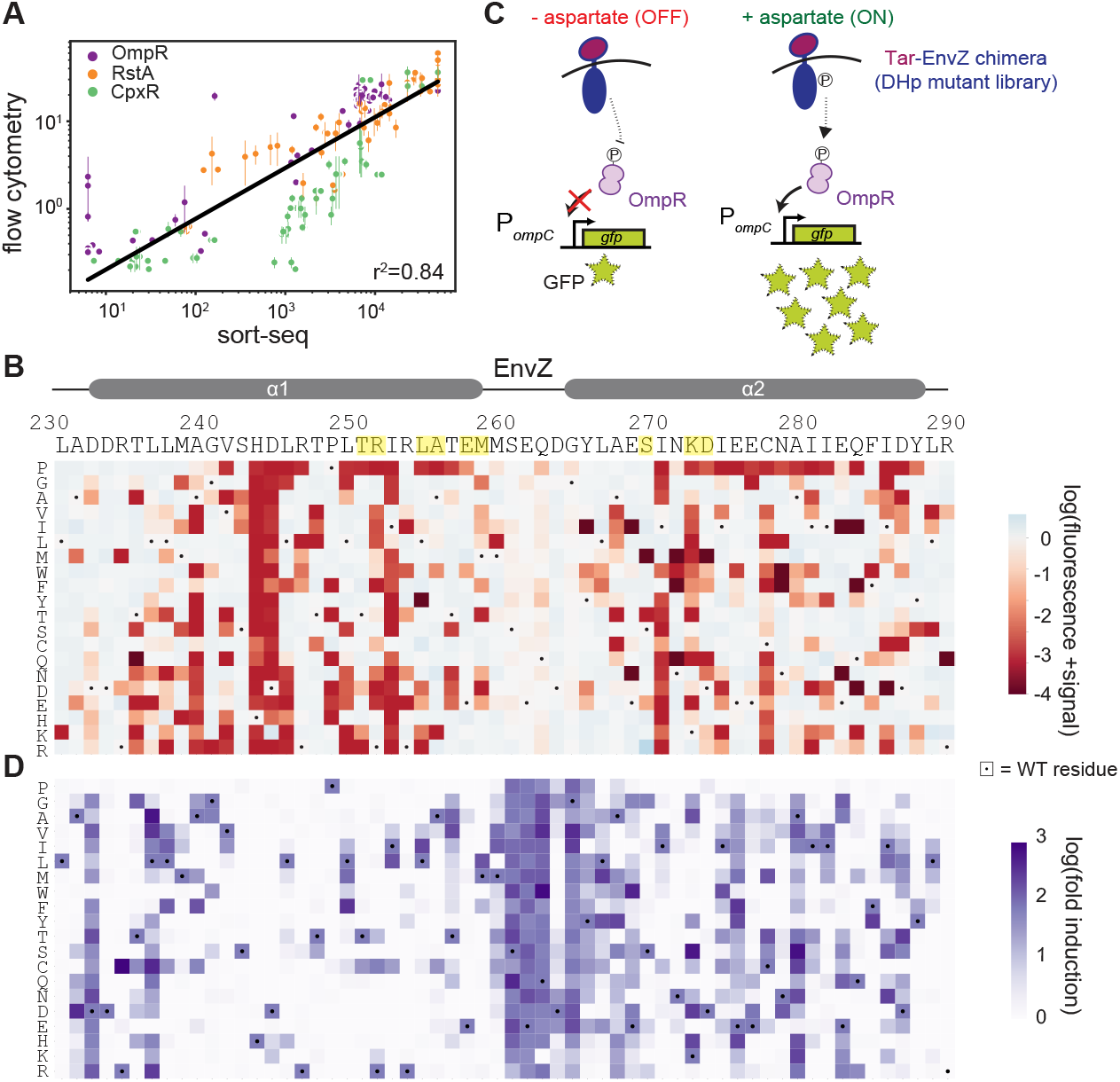
Sort-seq reveals the landscape of mutational tolerance in EnvZ-OmpR signaling. (A) Correlations between fluorescence values obtained by Sort-seq and by individual-clone flow cytometry. Error bars indicate S.D. from 2 independent biological replicates. Individual Pearson’s coefficients for the three reporters were r^2^ = 0.91 (OmpR), 0.92 (RstA) and 0.83 (CpxR). (B) Heatmap of OmpR reporter data with columns representing positions along the EnvZ DHp sequence; yellow highlights indicate coevolving residues. Rows indicate specific amino acids introduced at each position. Dots mark wild-type residues. Color-coded values represent log_10_(fluorescence) of each variant in the +signal condition. Wild-type EnvZ is set to white (blue represents increases in fluorescence, red shows decreases). (C) Diagram illustrating induction for the OmpR reporter. In low aspartate conditions, wild-type EnvZ is a phosphatase, removing phosphoryl groups from OmpR and leading to low GFP levels. In high aspartate conditions, wild-type EnvZ is a kinase, phosphorylating OmpR and driving high GFP production. (D) Same as (B) but with purple color indicating the log_10_(fold induction) value for each variant at each position.

### Deep mutational scanning reveals the marginal specificity of paralogs

To visualize our deep mutational scanning data for transfer to the cognate regulator OmpR, we generated a heatmap displaying the fluorescence levels in the presence of inducer (+signal) for each possible substitution at each position in the DHp domain of EnvZ (Fig. 2B). Most (81%) substitutions retained levels similar (within 5-fold) to wild-type EnvZ, indicating that they retain kinase activity (Fig. 2B). When visualizing fold-induction value (fluorescence +/- signal, Fig. 2C), 76% of substitutions eliminated or reduced the fold-induction relative to the wild-type (Fig. 2D). A clear exception was within the loop region connecting α1 and α2 where a wide range of substitutions was tolerated. The loss of signal responsiveness for many variants may result from reduced phosphatase activity in the absence of signal, producing a constitutively active kinase (Fig. S3B).

To quantify cross-talk from each EnvZ variant to CpxR, we generated a heatmap showing the increase in fluorescence of the CpxR reporter in the +signal condition relative to that of wild-type EnvZ, which was set to 0 (Fig. 3A). Increases in fluorescence represent increased kinase activity, which could disrupt signaling fidelity and constitute detrimental cross-talk (Fig. 3A, S4A–S4C). No single substitution produced an EnvZ variant with both kinase and phosphatase activity toward CpxR. Although the majority of substitutions in EnvZ did not increase cross-talk to CpxR, a small number of substitutions showed fluorescence values increased as much as 30-fold relative to wild-type EnvZ. This level of activation was similar to that of a chimera of the Tar sensor domain fused to the cytoplasmic signaling domains of CpxA, the cognate HK of CpxR (Fig. S4D). The cross-talk-inducing substitutions occurred primarily at the coevolving positions previously shown to be important for HK-RR interaction specificity^29,30^. For instance, at Ala255, Glu257, and Asp273, multiple substitutions with dissimilar biochemical characteristics caused cross-talk, suggesting that the native residues at these positions serve as negative design elements that prevent cross-talk to CpxR. At Ser269, only two substitutions, the positively charged residues Arg and Lys, caused substantial increases in cross-talk, suggesting that these residues may promote an interaction with CpxR that the wild-type Ser residue cannot. Notably, the corresponding residue of CpxA is Arg, consistent with positive charge at this position facilitating interaction with CpxR. At other coevolving positions, including Thr250, Arg251, Leu254, and Met258, no substitutions substantially increased cross-talk. These residues are each identical or biochemically similar in EnvZ and CpxA, consistent with them not being involved in insulating these two pathways.

**Fig. 3.**
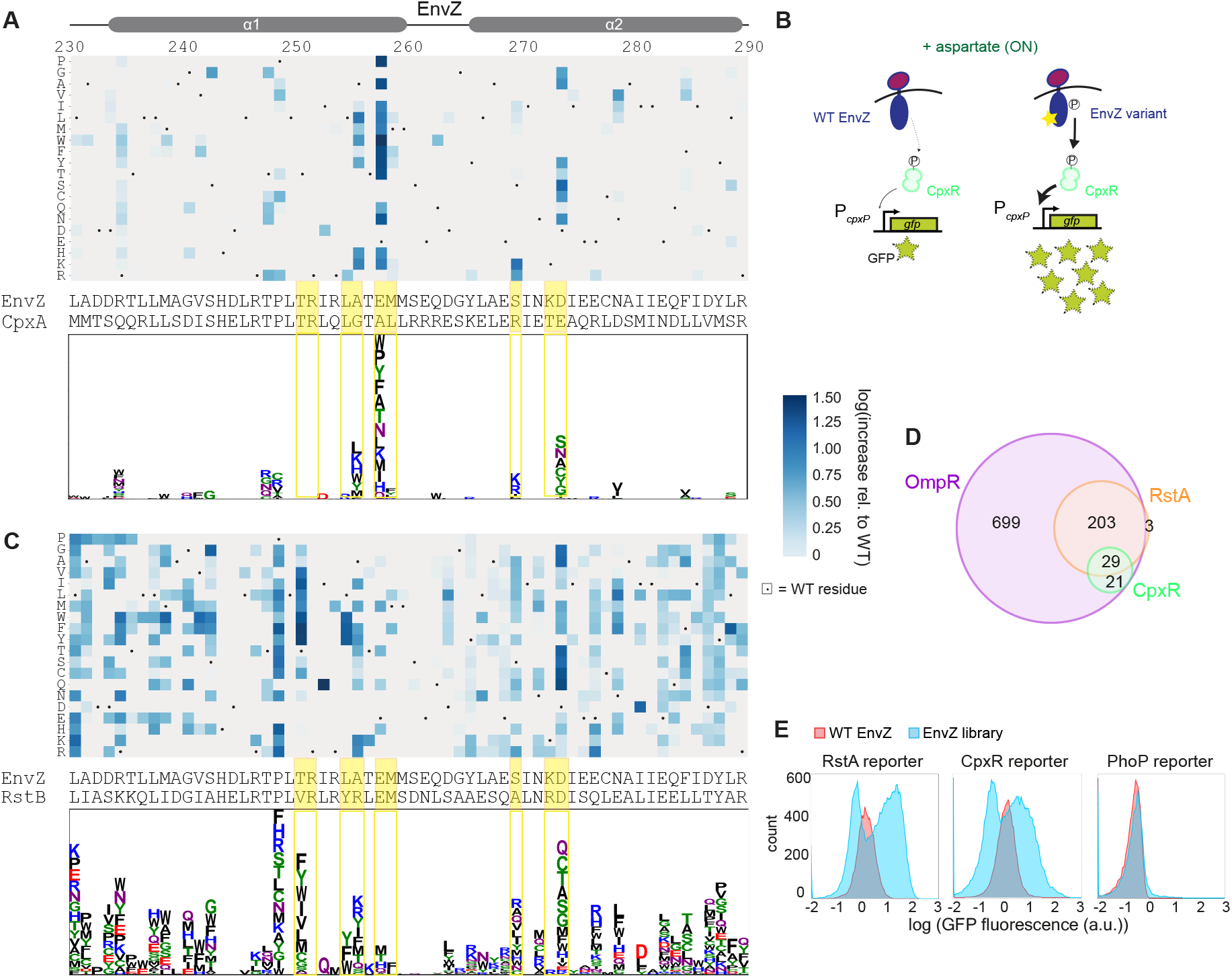
EnvZ exhibits marginal specificity, reflecting a crowded local sequence space. (A) Top: Heatmap of CpxR reporter data where values represent increases in GFP fluorescence relative to wild-type EnvZ in the +signal condition. Dots mark wild-type residues. EnvZ and CpxA DHp sequences are shown below. Yellow highlighted positions mark coevolving residues. Values were normalized by increases in fluorescence towards OmpR, which may reflect non-specific effects of a substitution on expression level or kinase activity that increase activity toward all RRs. Bottom: Logo transformation of heatmap where the height of a letter represents the increase in fluorescence relative to wild-type for that amino acid substitution at that position. Letters are stacked in ranked order. (B) Diagram illustrating fluorescence measurements for CpxR reporter. Wild-type EnvZ shows low levels of kinase activity, leading to low GFP levels. EnvZ variants (as indicated by the star) may show increased kinase activity, driving high GFP production. (C) Same as (A) except for RstA reporter data. (D) Overlap of single-substitution EnvZ variants with activity towards different RRs. The OmpR set contains variants with kinase activity for OmpR comparable to wild-type EnvZ (within 5-fold). The RstA and CpxR sets contain variants with ≥ 5-fold increases in activity towards RstA or CpxR relative to wild-type EnvZ. (E) Histograms of GFP fluorescence distributions for the RstA, CpxR, and PhoP reporters. Red populations are for wild-type EnvZ, blue populations are transformations of the single mutant library. Blue library populations for RstA and CpxR reporters are replicated from Fig. 1C for comparison to PhoP.

We also assessed cross-talk to RstA (Fig. 3C). The strongest cross-talk-inducing substitutions again tended to occur at the coevolving residues, with similar patterns seen as with CpxR. At some coevolving positions in which EnvZ differs significantly from the corresponding residue of RstB, such as Thr250, Ala255, Ser269, multiple biochemically distinct substitutions led to substantial cross-talk, indicating that these residues act as negative design elements with respect to RstA. At Leu254, only aromatic residues caused cross-talk suggesting that they form specific contacts with RstA that enhance its interaction with EnvZ; notably, RstB features a Tyr at this position. In a strikingly different pattern than we observed for CpxR, there were also a large number of substitutions at non-coevolving positions that caused cross-talk, which are discussed below.

In total, there were 21, 206, and 29 substitutions that increased cross-talk more than 5-fold toward CpxR, RstA, or both, respectively (Fig. 3D). Similar patterns and relative numbers of variants were seen at thresholds of 3- and 10-fold, indicating that our results are robust to the precise threshold used (Fig. S4E–S4G). The observation that many individual substitutions can readily produce cross-talk indicates that EnvZ exhibits marginal, rather than robust, specificity with respect to CpxR and RstA. Considering both CpxR and RstA, we found that EnvZ variants containing the corresponding residue of the respective cognate kinase (CpxA and RstB, respectively), were more likely to exhibit cross-talk relative to other substitutions (p = 0.0015, Fisher’s exact test; Fig. S5A). However, there were still a large number of EnvZ substitutions that did not resemble the corresponding residue on the other kinases but still caused cross-talk. Together, these findings demonstrate that mutating the EnvZ sequence to mimic RstB or CpxA is not the only way to generate cross-talk to RstA or CpxR (Fig. S5B).

We also found many substitutions that decreased the activity of EnvZ towards either or both non-cognate regulators, without decreasing activity toward the cognate regulator OmpR (Fig. S6A-C). We likely observed this only because EnvZ is overexpressed in our assay; at native levels, EnvZ shows no detectable activity towards RstA and CpxR (Fig. S6D and S6E). However, this finding does suggest that cross-talk between these pathways has not been eliminated, and instead has only been reduced to such a level that it has no effect on fitness. The notion that interactions with non-cognate proteins could be reduced further by many different single substitutions emphasizes that only marginal specificity has been selected for between these systems.

We hypothesized that the marginal specificity of EnvZ-OmpR, RstBA, and CpxAR reflects their phylogenetic history as closely related paralogs. Duplication events that led to the emergence of these three systems were likely followed by changes in specificity sufficient to insulate these pathways, such that they could carry out distinct functions, but leaving them close in sequence space. In contrast, less closely related pathways are likely further apart in sequence space such that specificity is more robust. To test this idea, we transformed the library of EnvZ variants into a reporter strain for the more distantly related regulator PhoP, for which the cognate kinase is PhoQ (Fig. S1B), and performed flow cytometry. Unlike with the RstA and CpxR reporter strains, there was no subpopulation of cells showing significantly increased fluorescence relative to wild-type EnvZ, indicating that no or very few single substitutions in EnvZ cause substantial cross-talk to the distantly related PhoP (Fig. 3E).

### The extent of marginal specificity reflects the evolutionary history of paralogs

Collectively, our results suggest that the occupancy of sequence space reflects the evolutionary history of paralogs and that cross-talk is most likely to occur between more closely related systems. Because EnvZ-OmpR is more closely related to RstBA than to CpxAR (Fig. S1B), this model may explain why many more single substitutions can cross-talk to RstA than CpxR (Fig. 3C). To assess the spatial distribution of substitutions that induced cross-talk, we mapped these substitutions onto a modeled structure of the EnvZ-OmpR complex in the phosphatase state, *i.e*., the state in which a HK is thought to promote dephosphorylation of its phosphorylated cognate RR (Fig. 4A). This is the state most commonly seen in two-component complex structures^39–41^, likely due to its rigidity facilitating crystallization. Substitutions that increased cross-talk to CpxR in our assay were generally on the surface of the HK dimer at the interface with the RR, which we refer to here as the primary interface. As noted above, these cross-talking substitutions largely involved residues known to coevolve in HK-RR pairs^29,30^ (Fig. 3A). In contrast, the substitutions that caused cross-talk to RstA mapped all over the DHp domain, including at positions distal to the interface, and even some within the core of the dimer (Fig. 4A).

**Fig. 4.**
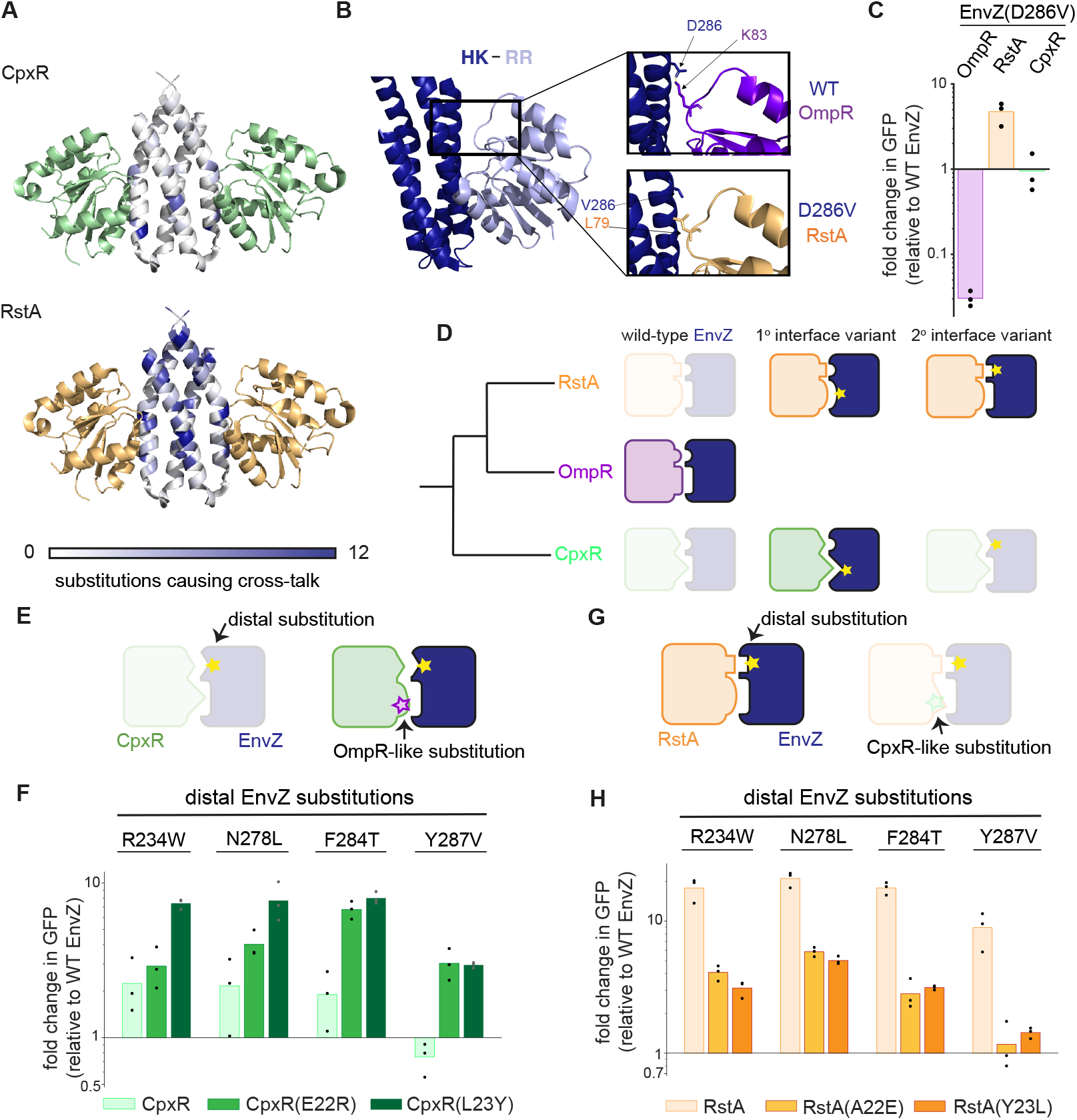
The degree of marginal specificity reflects phylogenetic relatedness between paralogs. (A) Number of substitutions causing ≥ 5-fold cross-talk to either CpxR (green) or RstA (orange) at each position in EnvZ is color-coded and mapped onto the model complex structure in the phosphatase state. (B) Active phosphotransfer state structure with HK in deep blue and RR in light blue (PDB: 5IUL). Insets show wildtype EnvZ and OmpR, or EnvZ(D286V) and RstA residues modeled onto this structure. Side chains were placed in the most preferred rotamers for interaction. (C) Fold change in fluorescence +signal for EnvZ(D286V) relative to wild-type EnvZ for each reporter strain. n=3 biological replicates. (D) Model in which faded protein pairs represent unsuccessful interaction and full-color pairs represent successful interaction. RstA and CpxR have diverged sufficiently from OmpR to prevent cross-talk with wild-type EnvZ. However, RstA retains enough compatibility that substitutions at either the primary or secondary interface (yellow stars) can produce cross-talk. By contrast, CpxR has diverged such that its primary interface is fundamentally incompatible with EnvZ, and only substitutions that suppress incompatibility at this interface create cross-talk. (E) Model in which distal substitutions do not create cross-talk to wild-type CpxR but substitutions at the CpxR primary interface that increase its compatibility with EnvZ can increase sensitivity to distal substitutions. (F) Fold changes in +signal fluorescence relative to wild-type EnvZ for four distal single substitutions in EnvZ, against wild-type CpxR and CpxR variants with OmpR-like primary interface substitutions: E22R and L23Y. n=3 biological replicates. (G) Model in which distal substitutions in EnvZ can produce cross-talk to RstA, but substitutions at the RstA primary interface that reduce its compatibility with EnvZ decrease its sensitivity to these distal substitutions. (H) Fold changes in +signal fluorescence relative to wild-type EnvZ for four distal single substitutions in EnvZ, against wild-type RstA and RstA variants with CpxR-like primary interface substitutions: A22E and Y23L. n=3 biological replicates.

We also modeled EnvZ-OmpR in the suspected kinase, or phosphotransfer, state, using the single complex that has been solved^42^. In this structure, the N- and C-terminal portions of the DHp domain, which form the upper part of the dimeric four-helix bundle, reside closer to the β4-α4 loop of the RR (Fig. 4B). Interactions at this secondary interface may explain the cross-talk behavior of some substitutions distal to the primary interface of both states. For example, the substitution D286V in EnvZ caused a ~50-fold decrease in activity towards OmpR, but a ~5-fold increase in activity towards RstA (Fig. 4C). Examining EnvZ-OmpR modelled onto the kinase-state structure, Asp286 is positioned such that it can form a salt bridge with Lys83 on the β4-α4 loop of OmpR (Fig. 4B). Substituting this Asp with a Val eliminates this favorable interaction, which likely explains the decreased transfer from EnvZ(D286V) to OmpR. The residue corresponding to Lys83 in RstA is Leu (L79), possibly explaining why a hydrophobic Val in place of Asp286 in EnvZ is more favorable for this interaction than the charged Asp (Fig. 4B).

Although additional interactions at the secondary interface may explain some of our results, they are unlikely to explain the effects of cross-talk-inducing substitutions within the DHp core. Such substitutions presumably impact the primary or secondary interface allosterically to enhance interaction with the non-cognate RstA. Importantly, almost no substitutions in the DHp core or at positions away from the primary interface created cross-talk with the less closely related CpxR (Fig. 4A). Thus, we hypothesized that the primary interfaces of EnvZ-OmpR and RstB-RstA have diverged enough to reduce cross-talk between wild-type EnvZ and RstA, but are still sufficiently compatible that a single substitution at a distal site can result in cross-talk. By contrast, the primary interface of CpxA-CpxR is far enough diverged from EnvZ-OmpR that only substitutions at this interface can yield large enough effects to produce cross-talk (Fig. 4D).

This hypothesis predicts that increasing the compatibility of the primary interface between EnvZ and CpxR should increase the propensity of distal substitutions in EnvZ to cause cross-talk (Fig. 4E). To test this idea, we sought to substitute interfacial residues in CpxR with those found in OmpR, and measure whether there is epistasis between these substitutions and distal substitutions in EnvZ. We focused on two coevolving interface positions at the primary interface of CpxR that differ significantly from the corresponding residues in both OmpR and RstA: Glu22 and Leu23 (Fig. S7A). We substituted each of these residues individually with the corresponding residue found in OmpR, making CpxR variants E22R and L23Y, and then tested these variants for interaction with wild-type EnvZ and a selection of EnvZ variants with substitutions (R234W, N278L, F284T, and Y287V) that are distal to the primary interface and caused cross-talk to RstA but not CpxR (Fig. S7B and S7C). For each EnvZ variant, there was significantly more cross-talk to the CpxR primary interface variants than to wildtype CpxR (Fig. 4F and S7D). This positive epistasis between interfacial residues of CpxR and interface-distal residues of EnvZ suggests that improved compatibility between EnvZ and CpxR at the primary interface increases the susceptibility of CpxR to cross-talk induced by single substitutions elsewhere in EnvZ.

Our model also predicts that decreasing the compatibility of the primary interface between EnvZ and RstA could have the opposite effect, decreasing the propensity of interface-distal substitutions in EnvZ to cause cross-talk (Fig. 4G). To test this prediction, we substituted the same primary interface positions in RstA to match those of CpxR, A22E and Y23L, and then tested these RstA variants for interaction with wild-type EnvZ and the same EnvZ variants as above. In each case, cross-talk to RstA caused by distal substitutions in EnvZ was significantly reduced for both RstA A22E and Y23L (Fig. 4H, S7E and S7F). This negative epistasis between interfacial residues of RstA and interface-distal residues of EnvZ further supports our model that the propensity for cross-talk between non-cognate proteins depends on latent compatibility between the proteins at the primary interface, which reflects the evolutionary history of the paralogs.

### Avoiding cross-talk is a pervasive selective pressure shaping sequence space occupancy

We conclude that EnvZ exhibits marginal specificity, reflecting a crowded local region of sequence space. This further suggests that the specificity of extant two-component signaling paralogs can be fragile, easily disrupted by single substitutions throughout the kinase. We sought to assess whether this marginal specificity has broadly affected EnvZ evolution.

First, we used HMMER to identify and align a set of 5,751 EnvZ orthologs from a wide range of proteobacteria. We then calculated the frequencies at which residues that caused cross-talk from *E. coli* EnvZ to either CpxR or RstA appear at the equivalent position in EnvZ orthologs from other species that also have RstBA and CpxAR. These cross-talk-inducing residues were found less frequently than residues that do not cause cross-talk (p = 1.2 x 10^-4^, D = 0.15, Kolmogorov-Smirnov test, Fig. 5A). We also found that a higher proportion of cross-talk-inducing residues were completely absent at the equivalent positions in EnvZ orthologs (p = 6.2 x 10^-4^, odds ratio = 0.15, Fisher’s exact test, Fig. S8A). These observations suggest that, even averaged across a large number of sequence backgrounds, the substitutions that we found to cause cross-talk may have been selected against, leading to their lower prevalence among EnvZ orthologs.

**Fig. 5.**
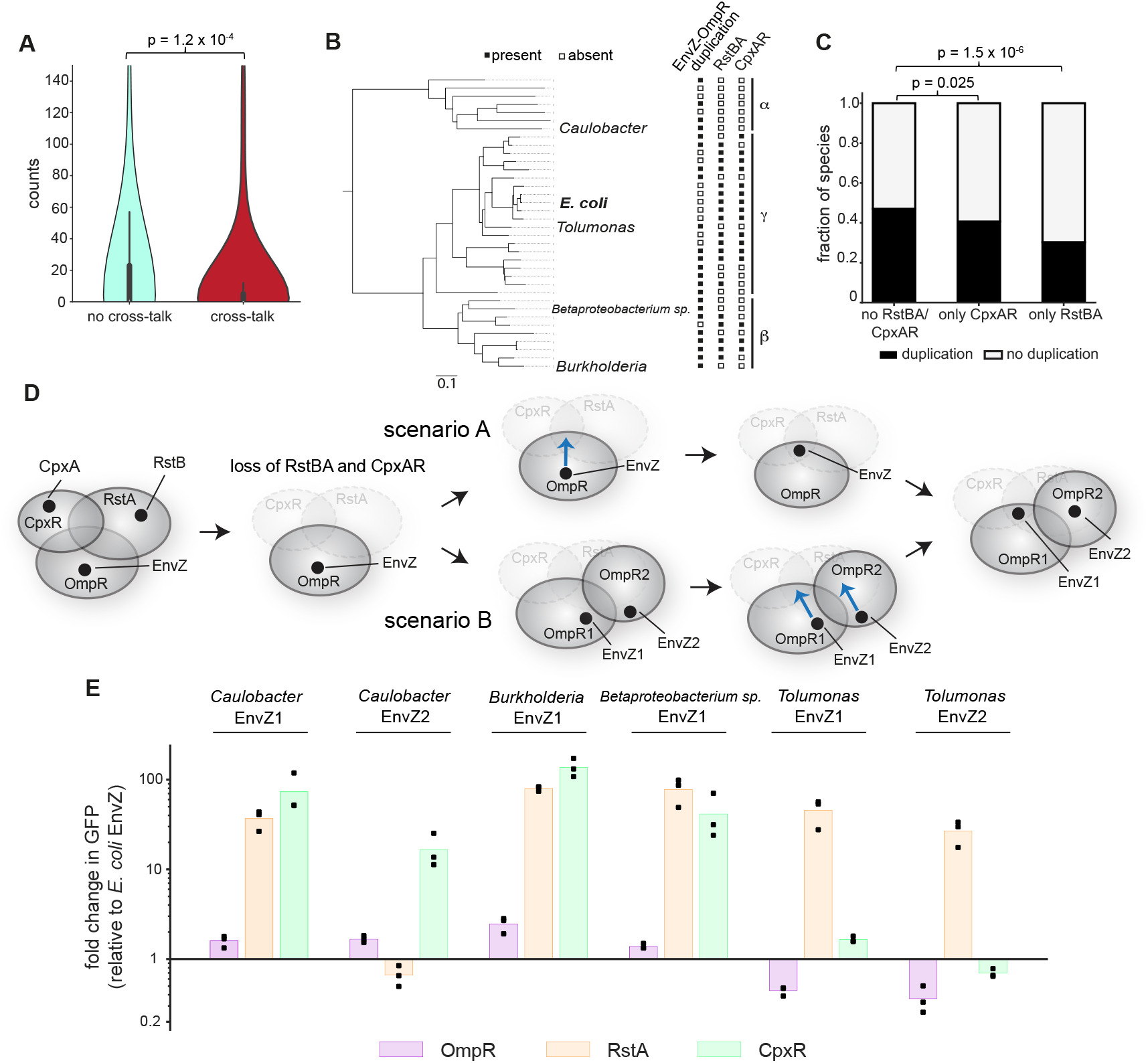
A densely occupied local sequence space constrains the evolution of two-component systems. (A) For single substitutions found to produce cross-talk or not to either RstA or CpxR (see Fig. 3), the violin plots show the distributions of counts of these substitutions found at the equivalent position in 1,019 EnvZ orthologs from species that also have RstBA and CpxAR (p = 1.2 x 10^-4^, Kolmogorov-Smirnov test). The inner box shows the quartiles and the whiskers show the range except for outliers. (B) Tree of a subset of species indicating whether they have RstBA and CpxAR and whether they have EnvZ-OmpR duplications. Closed and open squares indicate presence and absence, respectively. (C) Fraction of species that either do not have RstBA or CpxAR, have only CpxAR, or have only RstBA, that have duplications of EnvZ-OmpR (p = 0.025 for CpxAR loss, p = 1.5 x 10^-6^ for RstBA loss, Fisher’s exact test). (D) Sequence space diagram illustrating how species that have lost RstBA and CpxAR may relax the selection pressure against entering regions of sequence space that would previously have resulted in cross-talk, also freeing up sequence space for EnvZ-OmpR duplications. Blue arrow indicates a single mutation in EnvZ that would be tolerated only in species that have lost CpxR and RstA. This could occur before (scenario A) or after (scenario B) duplication. (E) Fold changes in fluorescence relative to *E. coli* EnvZ for six EnvZ homologs from four distantly related species, for *E. coli* OmpR, RstA, and CpxR reporters. n=3 biological replicates.

To test whether the differences seen were specific to EnvZ, we aligned ortholog sequences of three other HKs, PhoR, YehU, and BarA, which are increasingly distantly related to EnvZ (Fig. S1B). For YehU and BarA, there was no significant difference between the frequencies of the two classes of residues at the equivalent positions (p = 0.80, D = 0.043 for YehU, p = 0.18, D = 0.074 for BarA, Kolmogorov-Smirnov test, Fig. S8B and S8C) or the proportion of residues that were absent (p = 0.68, odds ratio = 0.94 for YehU, p = 0.17, odds ratio = 0.82 for BarA, Fisher’s exact test, Fig. S8D and S8E). For PhoR, there was a significant difference between the two categories, although it was smaller than the difference observed for EnvZ (p = 0.0016, D = 0.13, Kolmogorov-Smirnov test, p = 0.003, odds ratio = 0.12, Fisher’s exact test, Fig. S8F and S8G). These results suggest that more closely related kinases share some of the same sequence features and selective pressures faced by EnvZ, but these pressures do not apply to more distantly related kinases.

Although many γ-proteobacteria, like *E. coli*, have EnvZ-OmpR, RstBA, and CpxAR, several species have lost one or more of these systems (Fig. 5B). We wondered if losing RstBA or CpxAR relaxes the selection pressure against cross-talking mutations and allows drift of EnvZ into the regions of sequence space they previously occupied. Indeed, we found that cross-talk-inducing substitutions are seen at higher frequencies in EnvZ orthologs from species that have lost RstBA or CpxAR (p = 0.048, Kolmogorov-Smirnov test, Fig. S8H). Additionally, we found that species that have lost RstBA and CpxAR were more likely to have duplicated EnvZ-OmpR (p = 0.025 for CpxAR loss, p = 1.5 x 10^-6^ for RstBA loss, Fisher’s exact test, Fig. 5C). This finding suggests that the presence of these systems, particularly the most closely related RstBA, limits the sequence space available to EnvZ and thus constrains the ability of EnvZ-OmpR to duplicate and establish a new system (Fig. 5D).

We further predicted that in species lacking RstBA and CpxAR in which EnvZ-OmpR had duplicated, the EnvZ paralogs may now occupy sequence space made available by the loss of the other systems (Fig. 5D). To test this prediction we chose four species, distantly related to each other and to *E. coli*, in which RstBA and CpxAR were independently lost and EnvZ had been duplicated (Fig. 5B). Each of the resulting EnvZ homologs had residues that caused cross-talk in the context of *E. coli* EnvZ (Fig. S8I). These residues occurred at both primary and secondary interface positions, as well as in core residues of the DHp domain. We cloned and expressed each homolog in our reporter strains for *E. coli* OmpR, RstA, and CpxR, and then measured GFP expression relative to that seen with *E. coli* EnvZ. For each species considered, one or both EnvZ homologs showed high levels of cross-talk to *E. coli* RstA, CpxR, or both (Fig. 5E). This result strongly supports the notion that without RstBA and CpxAR, EnvZ duplicates commonly enter the sequence space freed up by the loss of these other systems. This finding further demonstrates how the presence of closely-related paralogs, and the consequent marginal specificity, has constrained EnvZ evolution.

## Discussion

Our findings demonstrate that the distribution of niches in sequence space of paralogous two-component signaling systems is not globally optimized for specificity or selected for robustness to mutation. Although the requirement for only a marginal level of specificity during the early establishment of duplicates may facilitate their evolution, it comes at the cost of future constraint on the emergence of additional duplications. Over time, due to drift and movement catalyzed by subsequent duplications, paralogous systems can continue to move apart in sequence space such that more distantly related systems are robustly insulated. However, systems like EnvZ-OmpR, CpxA-CpxR, and RstB-RstA, with ~2 billion years of divergence continue to exhibit marginal specificity. Thus, this drift is likely slower than the rate at which additional duplication events occur, such that the marginal specificity of existing paralogs will constrain the evolution of new duplicates when they emerge.

In other protein families, paralogs may share functional binding partners and not require all interactions to be fully specific. However, we expect that the principle of marginal specificity will apply in any case where there is a cost incurred by a non-specific interaction. Such cases are likely to occur between most paralogs whose functions are non-redundant.

Our results demonstrate how the nature of evolution, in only selecting for ‘good enough’, rather than fully optimized systems, results in small margins in specificity, and actually constrains the subsequent evolvability of paralogous families. This principle also applies to protein stability^21,43^, abundance^44^, and localization and assembly properties^45^. In each case, a large proportion of substitutions can disrupt the relevant property, suggesting that robustness has not evolved in these traits. The fragility of these properties has important implications for disease pathogenesis. For example, single substitutions that disrupt the stability and abundance of tumor suppressor proteins are implicated in cancer^44^, and single substitutions that affect assembly properties of proteins can drive hemoglobinopathies such as sickle cell anemia. The same appears to be true of specificity, where single ‘network-attacking’ substitutions that alter the specificity profile of human kinases are thought to disrupt cellular signaling networks and contribute to cancer progression^46^.

In addition to shedding light on the fundamental mechanisms of evolution and their consequences for paralogous families, our findings also have implications for protein design and directed evolution methods. Attempts to build new signaling systems while avoiding detrimental cross-talk with existing cellular systems may work better if employing randomization or mutagenesis of multiple residues, allowing ‘jumps’ into sparsely occupied regions of sequence space, rather than methods that traverse crowded local sequence spaces by moving one mutation at a time. Overall, we demonstrate an example of marginal specificity in protein interactions that has implications for the evolvability of paralogous families, in both natural and synthetic settings.

## Methods

### Bacterial strains and media

*E. coli* strains were grown in M9 medium (1× M9 salts, 100 μM CaCl_2_, 0.2% glucose, 2 mM MgSO_4_, with or without 5 mM aspartate). When indicated, antibiotics were used at the following concentrations: carbenicillin 50 μg/ml, kanamycin 50 μg/ml, spectinomycin 50 μg/ml and chloramphenicol 32 μg/ml. The base strain for all studies was *E. coli* strain ML1803 (Yale BW28357 *DenvZ*, Table S1)^32^. The OmpR reporter strain contained a p15a/cmR plasmid containing P_*ompC*_-*gfp*^32^ (Table S2). The RstA, CpxR, and PhoP reporter strains each contained additional deletions: *DackA-pta* (removes a pathway that generates acetyl phosphate, which can phosphorylate RRs in the absence of a HK) and *DrstB, DcpxA*, or *DphoQ* (to prevent the cognate HKs of RstA, CpxR, or PhoP, respectively, from phosphorylating or dephosphorylating them, Supp. Table 1), and the same reporter plasmid but with P_*asr*_-P_*cpxP*_-*gfp* or P*_mgrB_-gfp*, respectively (Supp. Table 2).

All libraries were cloned onto a low-copy pSC101/spec^R^ plasmid in which Taz variant expression was driven by a constitutive P_*lpp*_ promoter (Supp. Table 2). Characterization of individual, specific Taz variants were done using the same plasmid. EnvZ point mutations or homolog sequences were introduced using Gibson assembly using primers DG001-066 (Supp. Table 3). For the experiments involving mutations in the RRs, genomic mutations were made. Deletions discussed above and genomic mutations were made using *sacB-kan^R^* cloning. The loci in the relevant reporter strains were first replaced with the *sacB-kan^R^* locus using recombination^47^, and selected using kanamycin resistance. The *sacB-kan^R^* loci were then replaced using DNA fragments that corresponded to either the sequence of clean deletions, or genes with the relevant mutations, and selected using negative selection on sucrose. The relevant region of the genome was amplified by PCR and sequenced to confirm that the deletions/mutations were correct.

### Flow cytometry characterization

To induce Taz, cells were grown to early exponential phase (optical density at 600 nm (OD600) of about 0.2) in M9 before adding aspartate to a final concentration of 5 mM. Cells were grown for 3 h, diluted 1:40 into PBS with 0.5 g/L kanamycin, and fluorescence was measured on a Miltenyi MACSQuant VYB. In each cytometry experiment, three colonies of each strain were grown and induced independently and 30,000 cells were measured per replicate. FlowJo was used to analyze the data, gating on single live cells and extracting the median of the GFP distribution.

### Design and assembly of the Taz library

A comprehensive single mutant library was constructed using oligonucleotide-directed mutagenesis of the EnvZ DHp domain. To mutate each position in the Taz DHp (positions 230–289), two complementary 30-nucleotide primers (one sense, one antisense) were synthesized that introduce an NNS codon at the targeted position (primers DG067-186, Supp. Table 3). N is a mixture of A, T, C and G, and S is a mixture of G and C. This mutagenesis strategy results in 32 possible codons, which cover all 20 amino acids. One round of PCR was carried out with one reaction containing the antisense NNS primer and DG187, a primer located 105 bp upstream of the 5’ end of the DHp domain, containing a SacI restriction site, and a second reaction containing the sense NNS primer and DG188, a primer flanking the 3’ end of the *taz* gene, containing a SalI restriction site. A second PCR round using both first round products and both flanking primers produced the full-length double stranded product. All reactions yielded a band of the correct size on an agarose gel, which was extracted and purified (Zymo). PCR product concentrations were quantified (NanoDrop), pooled in equimolar ratios, digested with SacI and SalI, and ligated into the pSC101/spec^R^ expression vector. Each ligation was dialyzed on Millipore VSWP 0.025-μm membrane filters for 60 min and the entire volume was electroporated into 20 μL Invitrogen One Shot TOP10 Electrocomp *E. coli* to yield ~10^6^ total transformants. Plasmids from these transformants were then purified by miniprep (Zymo), dialyzed, and electroporated into the experimental strains, yielding ~10^9^ transformants for each strain.

### Sort-seq

For each of three replicates, 1 mL of overnight culture of the library was washed with M9 and inoculated into 50 mL of M9. Cells were grown to OD600 = 0.2 and each culture split into two: aspartate was added to a final concentration of 5 mM to one of the cultures. After 3 h, cells were diluted (1:30) into PBS containing 320 μg/ml chloramphenicol and cells were placed on ice for sorting. Cells were sorted into bins based on GFP expression on a BD FACS Aria machine. Single live cells were isolated using the gating strategy in Fig. S2J. The FITC voltage was adjusted so that the population spanned the range of fluorescence the machine could detect. A live histogram of FITC fluorescence was drawn and gates were spaced evenly along the log10(GFP) axis. For each library replicate, both the on and off cultures were sorted into 8 separate bins, generating 48 total bins. Up to ~2 million cells were sorted into bins per replicate (Fig. S2G-I). Sorted cells were added to 2× YT medium containing chloramphenicol and spectinomycin, and then grown overnight.

### Illumina sample preparation

After FACS, plasmids were purified (Zymo) from overnight cultures representing each bin from each library replicate. Two PCR reactions were performed, both using KAPA HiFi, to add Illumina sequencing adaptors and barcodes. First, DHp domain sequences were amplified for 12 cycles (95 °C for 30 s, 68 °C for 15 s and 72 °C for 30 s) with Illumina inner amplification primers (primers DG189-198, Table S3). Second, purified PCR product from the first reaction was amplified in a second PCR with barcoding primers (primers DG199-224, Supp. Table 3) for 9 cycles (95 °C for 30 s, 68 °C for 15 s and 72 °C for 30 s). Final products were quantified (NanoDrop), normalized, combined, and sequenced on an Illumina NextSeq. For each bin, 1–2 million reads were collected.

### Illumina data processing

Sort-seq data processing was carried out as previously described^9^. The frequency of each variant in each bin was calculated by taking the fraction of reads in a given bin that corresponded to a given sequence, normalized by the fraction of cells in that given bin. For each variant, the mean frequencies in each bin across three replicates, and standard deviation were used to fit Gaussian functions to each distribution (in log_10_(GFP units)), from both the on and off sorts (SciPy optimize package). Fold-induction values were calculated as the ratio of the means between the induced and uninduced states: μon/μoff. To assess cross-talk to CpxR and RstA, fluorescence values in the presence of inducer were normalized against increases in fluorescence towards OmpR, which may reflect non-specific effects of a substitution on expression level or kinase activity that increase activity toward all three RRs. Gaussian fit means for each variant for reporter can be found in Table S4 (OmpR), Table S5 (RstA) and Table S6 (CpxR).

### Purification of two-component signaling proteins and in vitro phosphotransfer assays

Expression and purification of EnvZ variants and RRs, and phosphotransfer experiments, were carried out as previously described^29,30,48^. RRs were fused to a His_6_–Trx tag, and the cytoplasmic region of EnvZ (residues 222–451) was fused to a His_6_–MBP (maltose-binding protein) tag, expressed in BL21(DE3) cells and purified on a Ni^2+^-NTA column. For phosphotransfer reactions, the kinase was autophosphorylated for 1 h at 30 °C with [γ-^32^P]ATP (Perkin Elmer) before being combined with RRs at a 1:4 ratio (10-μl reactions contained 1 μM EnvZ and 4 μM RR). Reactions were stopped at the times noted by adding 4× Laemmli buffer with 8% 2-mercaptoethanol. HKs and RRs were separated by SDS-PAGE, gels were incubated with phosphor screens and imaged using a Typhoon imager (GE Healthcare) at 50 mm resolution. A representative image of two independent experiments is shown in Fig. S3A.

### Identification and assembly of ortholog sequences and trees

A tree of two-component systems was built by constructing a Hidden Markov Model profile from an alignment of DHp and CA domains of *E. coli* histidine kinases^29^ and searching the ProGenomes2.0 database^49^ with this profile. The phylogenetic tree shown in Fig. S1C was constructed using FastTree and pruned to display only *E. coli* systems. EnvZ, RstB, and CpxA homologs from the ProGenomes2.0 database were identified and aligned using HMMER. A phylogenetic tree was constructed using FastTree and orthologs were classified based on clade identity. The resulting collection of sequences was further filtered by reciprocal HMMER to confirm that the best hit for each sequence in the *E. coli* genome was the correct paralog out of EnvZ, RstB, and CpxA. Each sequence maintained its species ID, allowing species with or without the relevant paralogs to be identified. The same process allowed YehU, BarA, and PhoR homologs to be identified, aligned and filtered. Species trees in Fig. S1B and Fig. 5B were obtained by pruning the tree constructed in ref^48^. Progenomes 2.0 species IDs for species tested in Fig. 5E are 155892 (Caulobacter), 1770053 (Burkholderia), 1797492 (Betaproteobacterium sp.), 595494 (Tolumonas).

### Statistical calculations

Two-sided Kolmogorov-Smirnov tests were used to determine whether there was a significant difference in distribution between counts of substitutions that do or do not cross-talk in MSAs of HK orthologs. Two-sided Fisher exact tests were used to determine whether there was a significant difference between presence/absence within the MSA of substitutions that do or do not cross-talk. Enrichment in Fig. S8H is calculated as counts in the first category of species, with counts in the second category subtracted, after scaling for the number of species in each category.

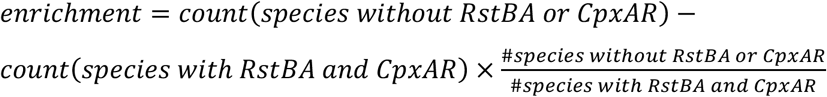

## Data availability

Datasets generated during this study are available at NCBI SRA to reviewers at https://dataview.ncbi.nlm.nih.gov/object/PRJNA902002?reviewer=vlkung36sasi433al3ou6vbin0. Datasets will be released to the public upon publication.

## Code availability

Python scripts for analysis are available at https://github.com/d-ghose/laub.

## Acknowledgements

We thank I. Nocedal for help with bioinformatic code and datatsets. We thank S. Srikant, I. Frumkin, D. Saxton, and S. Swanson for helpful discussions. M.T.L. is an Investigator of the Howard Hughes Medical Institute. This work was also supported by AFO MURI FA9550-22-1-0316 to M.T.L.

## Contributions

D.A.G., A.E.K. and M.T.L. conceptualized and designed the study. D.A.G. performed all experiments with assistance from K.E.P. and E.M.M. D.A.G., M.T.L., and A.E.K. wrote the manuscript. A.E.K. and M.T.L. supervised the study and provided funding support.

## Supplemental Figures

**Fig. S1.**
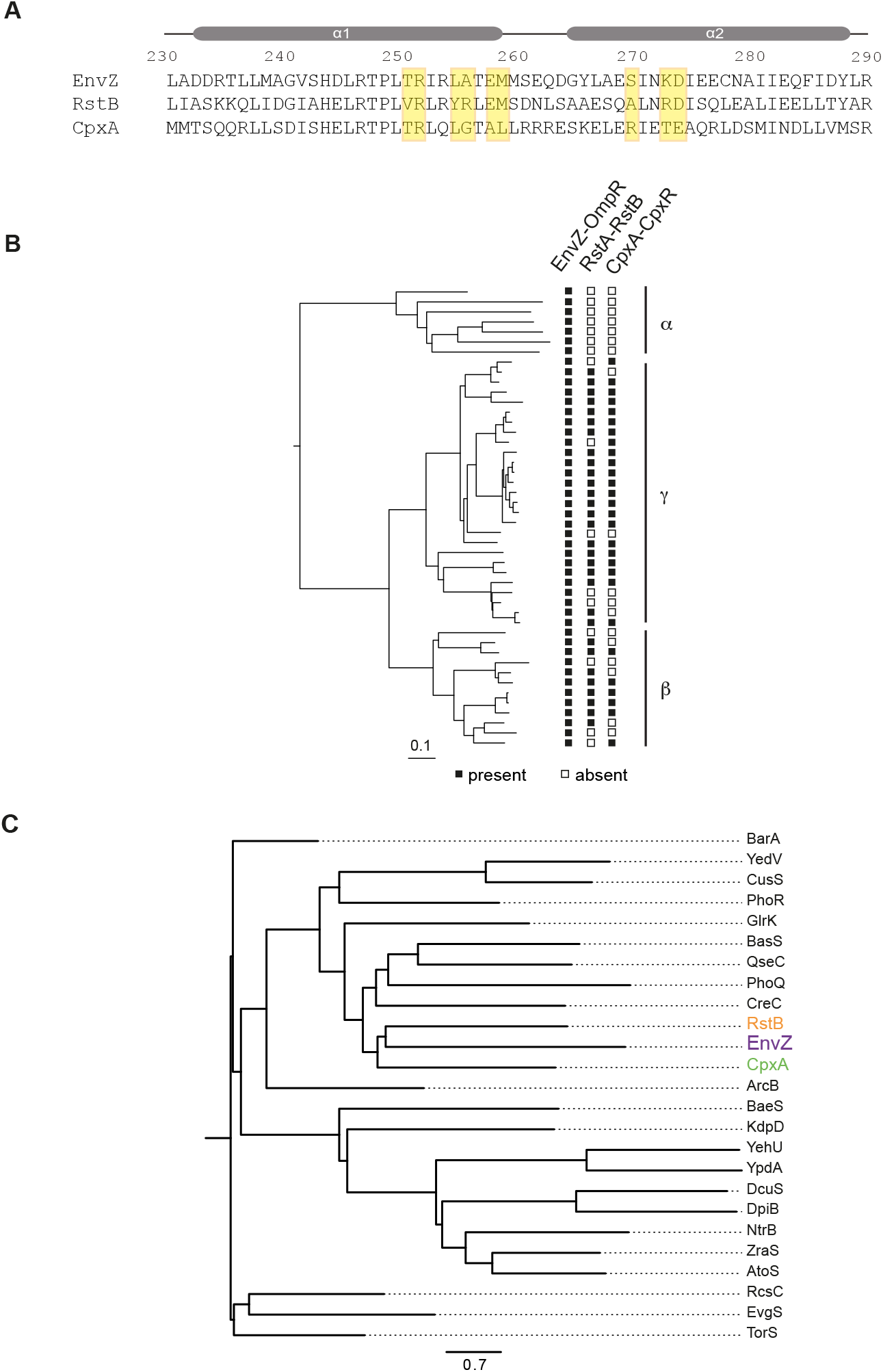
Phylogenetic analysis of *E. coli* two-component signaling systems. (A) Alignment of DHp domain sequences from EnvZ, RstB, and CpxA. Residues that strongly coevolve in HK and RR proteins are highlighted in yellow. (B) Phylogenetic tree of α-, β-, and γ-proteobacteria, where closed and open squares indicate presence and absence, respectively, of EnvZ-OmpR, RstBA, and CpxAR. Scale bar indicates substitutions per site. (C) Tree of *E. coli* histidine kinases with systems investigated in this study in larger text and color. Scale bar indicates substitutions per site.

**Fig. S2.**
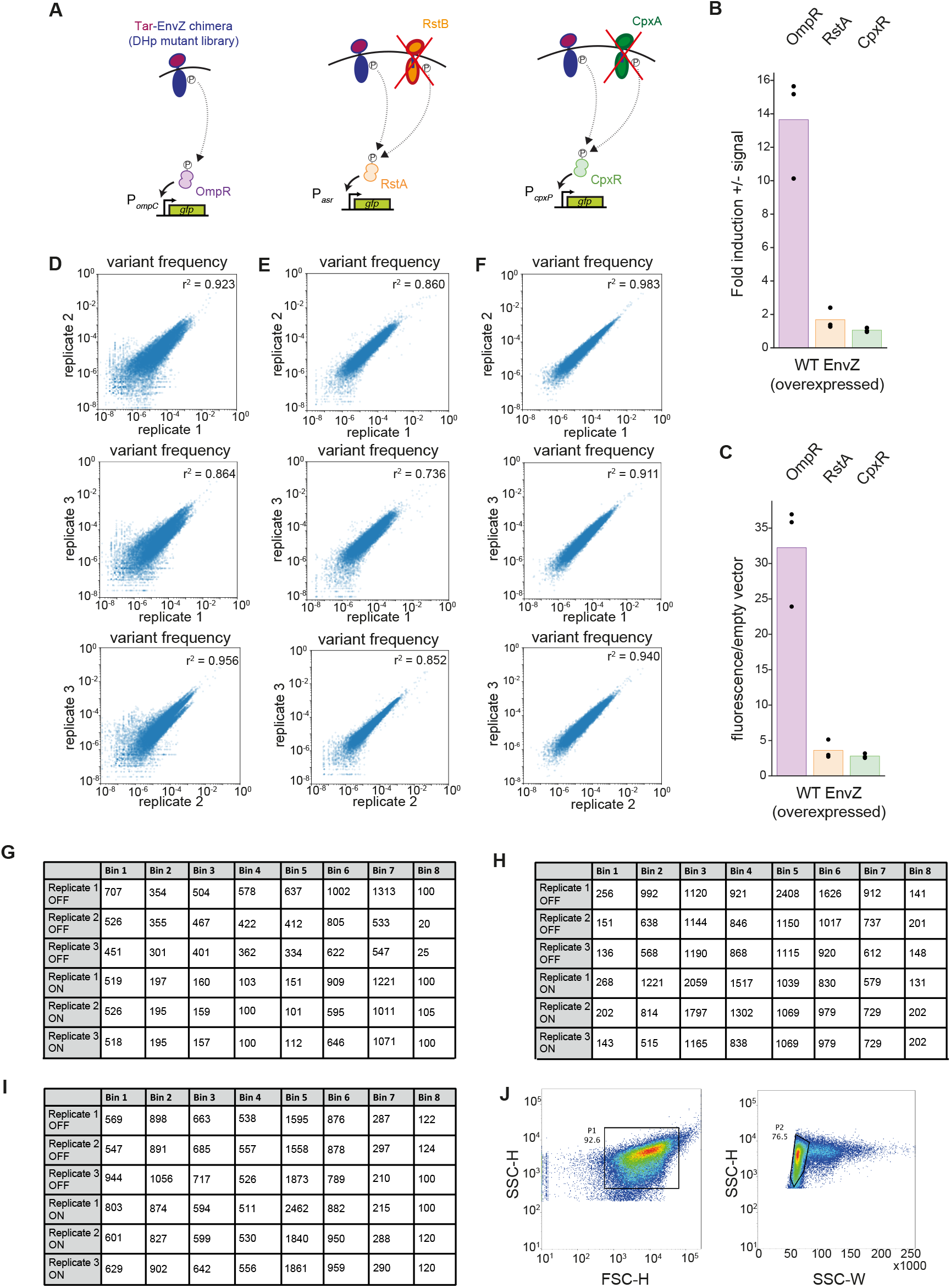
Details of Sort-seq method and summary statistics. (A) Summary of the three reporter strains used: OmpR reporter plasmid in *DenvZ* strain, RstA reporter plasmid in *DenvZ DrstB Dacka-pta* strain, CpxR reporter plasmid in *DenvZ DcpxA Dacka-pta* strain. Cognate HKs of RstA and CpxR (RstB and CpxA) were removed from their reporter strains to prevent phosphorylation or dephosphorylation by these kinases from affecting the readout. (B) Fold induction (+/- signal) of each reporter with wild-type EnvZ expressed at the level used in the library. n=3 biological replicates. (C) Fluorescence levels (+ signal) of each reporter with wild-type EnvZ expressed at the level used in the library. Levels were normalized to an empty vector control for each strain. n=3 biological replicates. (D-F) Scatter plots displaying the correlations between the bin frequencies of individual variants measured in independent replicates for the OmpR (D), RstA (E), and CpxR (F) reporters. (G-I) Counts of cells sorted into each bin, in thousands, for each replicate and condition, for the OmpR (G), RstA (H), and CpxR (I) reporters. (J) FACS gating strategy for isolating live single cells. SSC-H/W = Side Scatter Pulse Height/Width, FSC-H = Forward Scatter Pulse-Height.

**Fig. S3.**
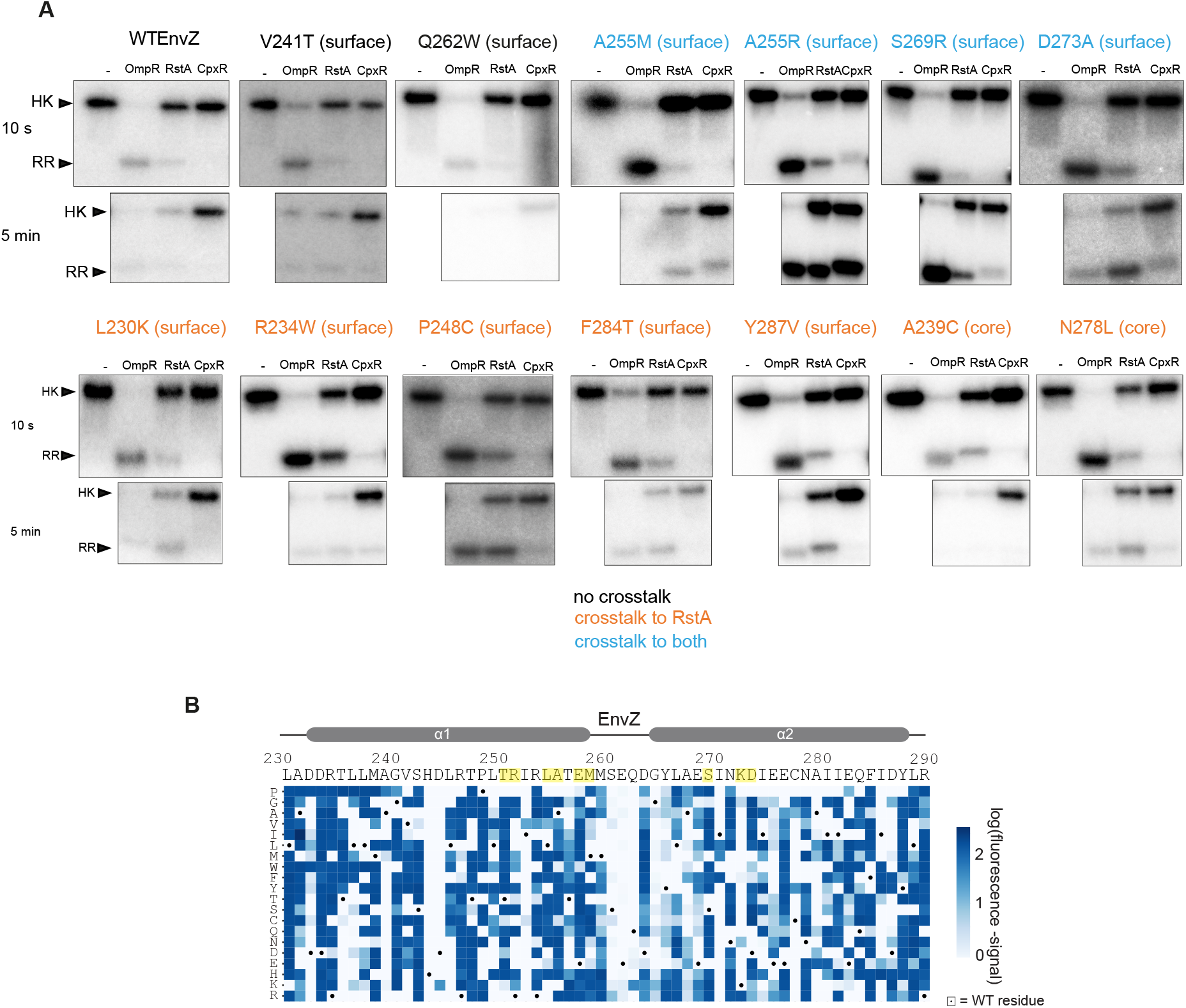
Validation of mutation effects and additional OmpR reporter analysis. (A) Phosphotransfer assays conducted with a range of EnvZ variants against OmpR, RstA, and CpxR at 10 s and 5 min time points. The upper band shows ^32^P-ATP incorporated into the autophosphorylated kinase, which depletes upon phosphotransfer to the response regulator. The lower band shows ^32^P-ATP incorporated into the response regulator, which at first accumulates upon transfer, and then depletes due to phosphatase activity by the kinase. Color of the text listing a given substitution indicates cross-talk activity seen in the deep mutational scanning, with black, orange, and blue indicating no cross-talk, cross-talk to RstA, and cross-talk to both, respectively. Wild-type EnvZ and two variants that showed no cross-talk activity in the screen are shown, along with 4 substitutions at coevolving residues that showed cross-talk to RstA and CpxR, and 7 substitutions at distal positions that showed cross-talk to RstA. Residues on the surface or within the core of the DHp domain are indicated. (B) Heatmap of OmpR reporter data with columns representing positions along the EnvZ DHp sequence; yellow highlights indicate coevolving residues. Rows indicate specific amino acids introduced at each position. Dots mark wild-type residues. Blue color indicates the log_10_(fluorescence) value of each variant in the -signal condition. Wildtype EnvZ is set to white (blue represents increases in fluorescence). Many variants have increased fluorescence in this condition suggesting that they have reduced phosphatase activity and are constitutively active.

**Fig. S4.**
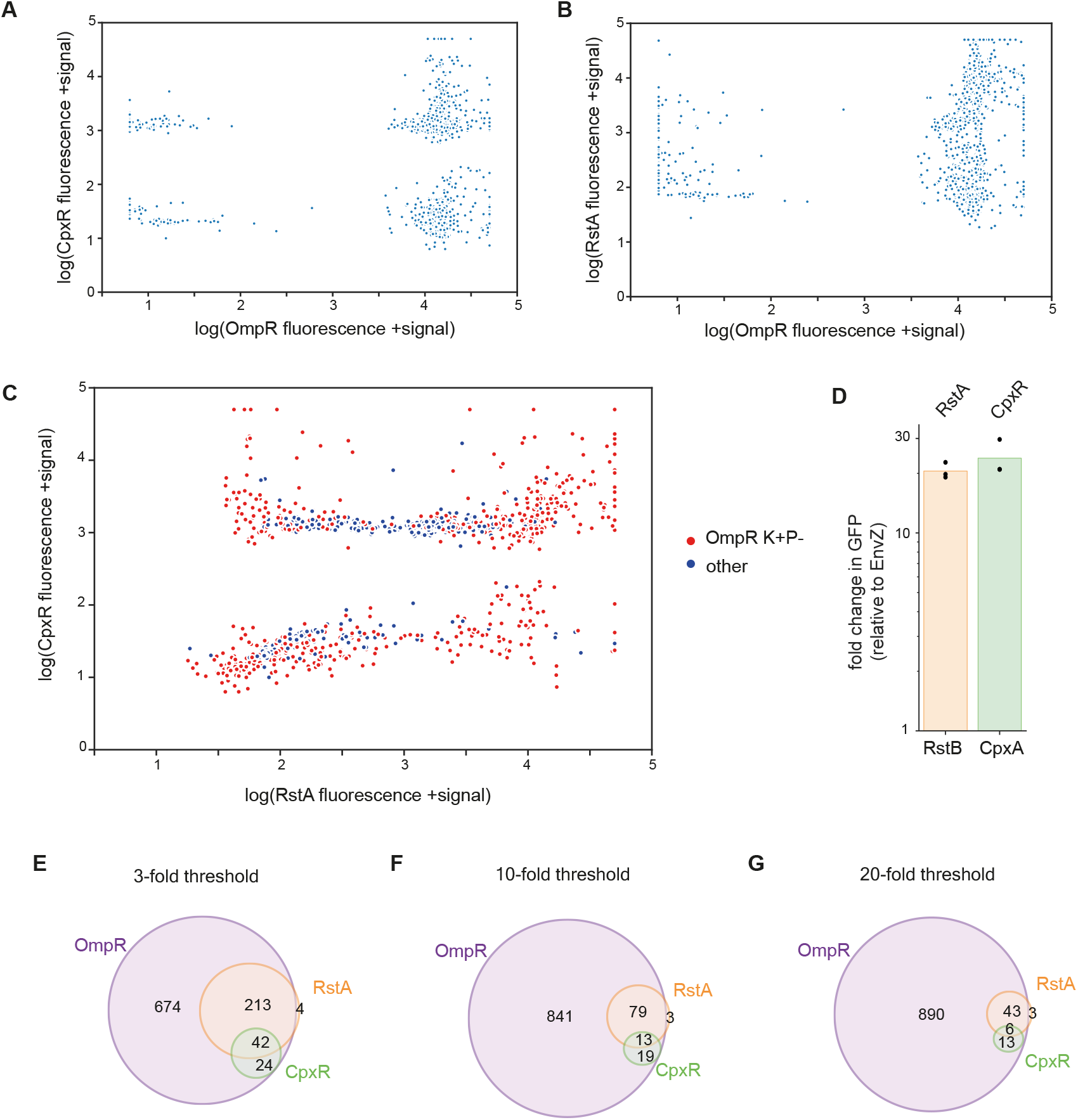
Validation of mutation effect sizes and thresholds. (A-B) Scatter plot displaying the correlations between fluorescence values of EnvZ variants screened against the (A) CpxR and OmpR reporters or (B) RstA and OmpR reporters in +signal condition. (C) Scatter plot displaying the correlations between +signal fluorescence values of EnvZ variants screened against CpxR vs. RstA. Red dots show variants that retain kinase activity but not phosphatase activity towards OmpR (K+P-variants). There is little correlation between the fluorescence values of variants with the three different reporters, showing that mutation effects are largely specific to each RR, and not caused by general properties such as expression level or kinase activity. There is also no relationship between loss of phosphatase activity towards OmpR and mutation effect towards RstA and CpxR, demonstrating that simply perturbing the equilibrium of kinase/phosphatase activity towards kinase is not sufficient to generate cross-talk. (D) Fold change in fluorescence of RstA and CpxR reporters relative to wild-type EnvZ when tested with chimeric HKs containing the Tar sensor domain and RstB or CpxA cytoplasmic domains, respectively. n=3 biological replicates. (E-G) Overlap of single-substitution EnvZ variants with activity towards different RRs. The OmpR set contains sequences which showed kinase activity towards OmpR at a comparable level to wild-type EnvZ (within 5-fold). The RstA and CpxR sets contain sequences which showed (E) ≥ 3-fold, (F) ≥ 10-fold, or (G) ≥ 20-fold increases in activity towards RstA or CpxR relative to wild-type EnvZ.

**Fig. S5.**
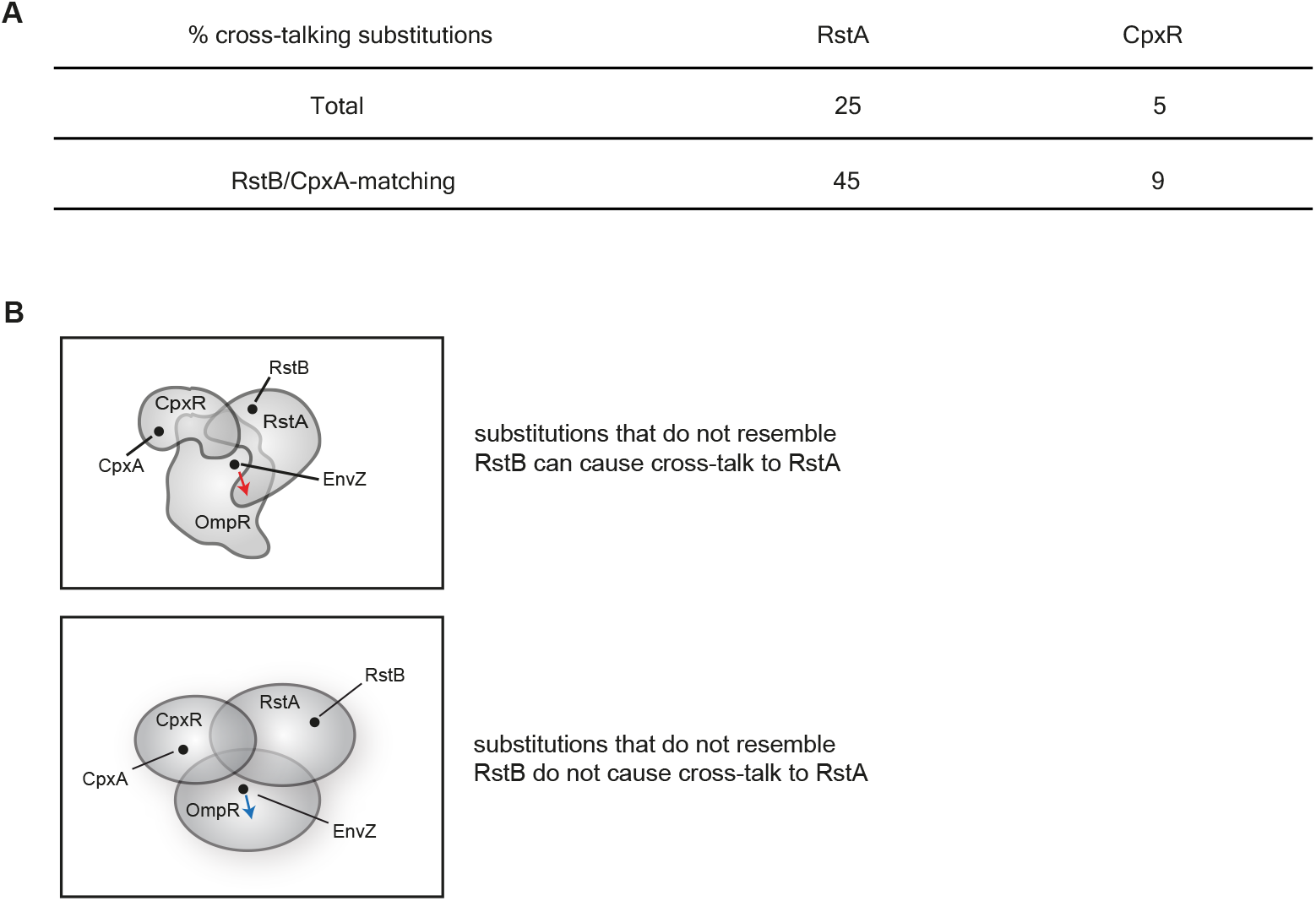
Niches in sequence space are irregularly shaped. (A) Table showing percentage of functional sequences (that have kinase activity for at least one regulator) that crosstalk to RstA or CpxR, among all functional sequences (top row) or among functional sequences that have a substitution corresponding to the residue found at the equivalent position in the cognate HK of RstA or CpxR (bottom row). (B) Sequence space diagram illustrating the irregular shapes of two-component signaling protein niches. Changing the EnvZ sequence to be more similar to RstB or CpxA is not the only way to generate cross-talk towards RstA or CpxR.

**Fig. S6.**
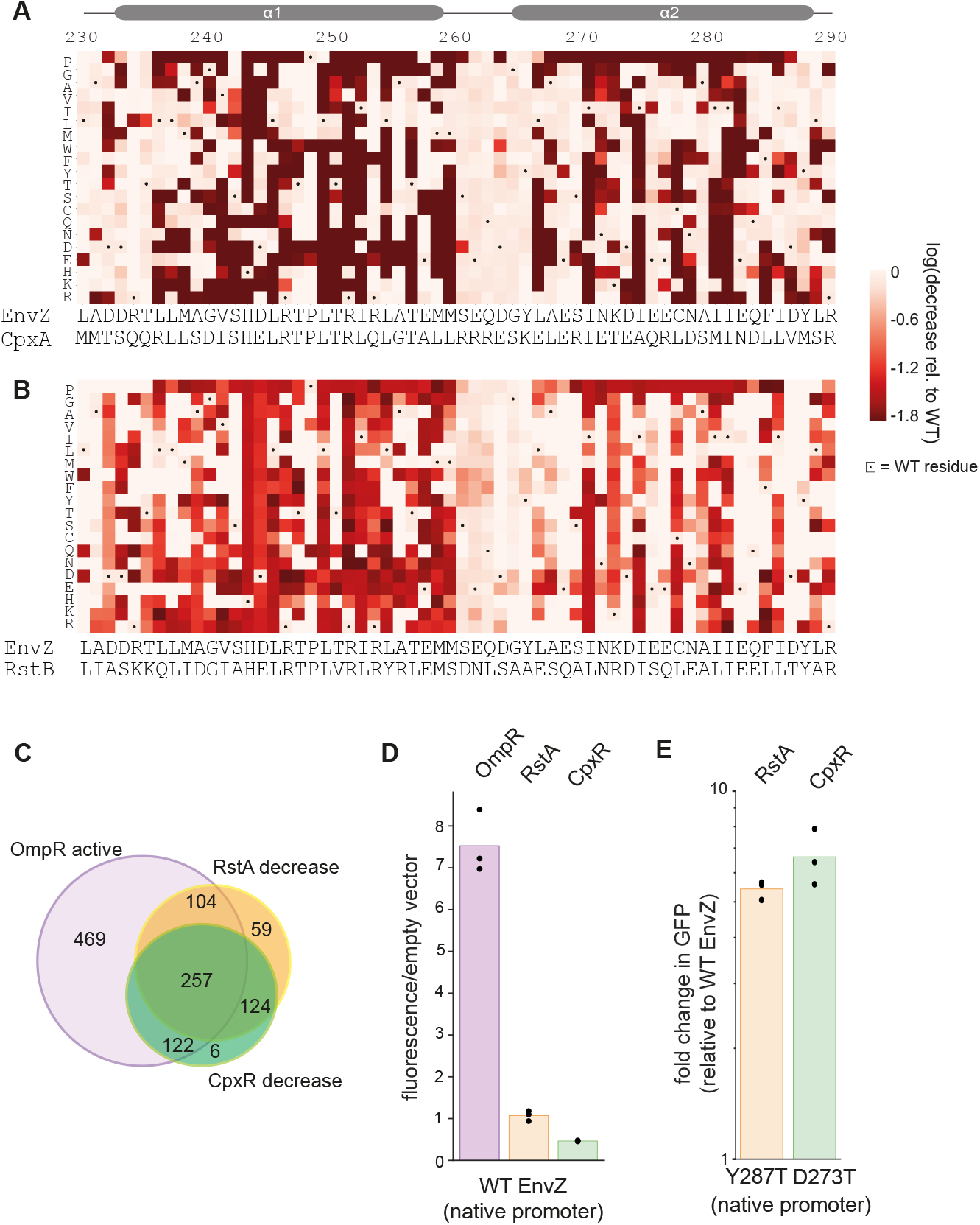
Many substitutions reduce interactions with non-cognate RRs. (A-B) Heatmaps of (A) CpxR or (B) RstA reporter data where the values represent log10(fluorescence) of variants in +signal condition. Wild-type EnvZ is set to 0, and color-coded as white, and is the maximum value shown (red shows decreases). All variants with increased fluorescence are shown as 0. Dots mark wild-type EnvZ residues. (C) Overlap of single-substitution EnvZ variants with activity towards different RRs: the OmpR set contains sequences that show kinase activity towards OmpR at a comparable level to wild-type EnvZ (within 5-fold). RstA and CpxR sets contain sequences with ≥ 5-fold decreases in activity towards RstA or CpxR relative to wild-type EnvZ. (D) Raw fluorescence levels for OmpR, RstA, and CpxR reporters following expression of wild-type EnvZ under its native promoter, normalized to levels seen with an empty vector for each strain. n=3 biological replicates. (E) Variants that show 5-fold cross-talk to RstA or CpxR in the Sort-seq screen (where kinase variants are overexpressed) still show cross-talk when expressed from the native promoter. n=3 biological replicates.

**Fig. S7.**
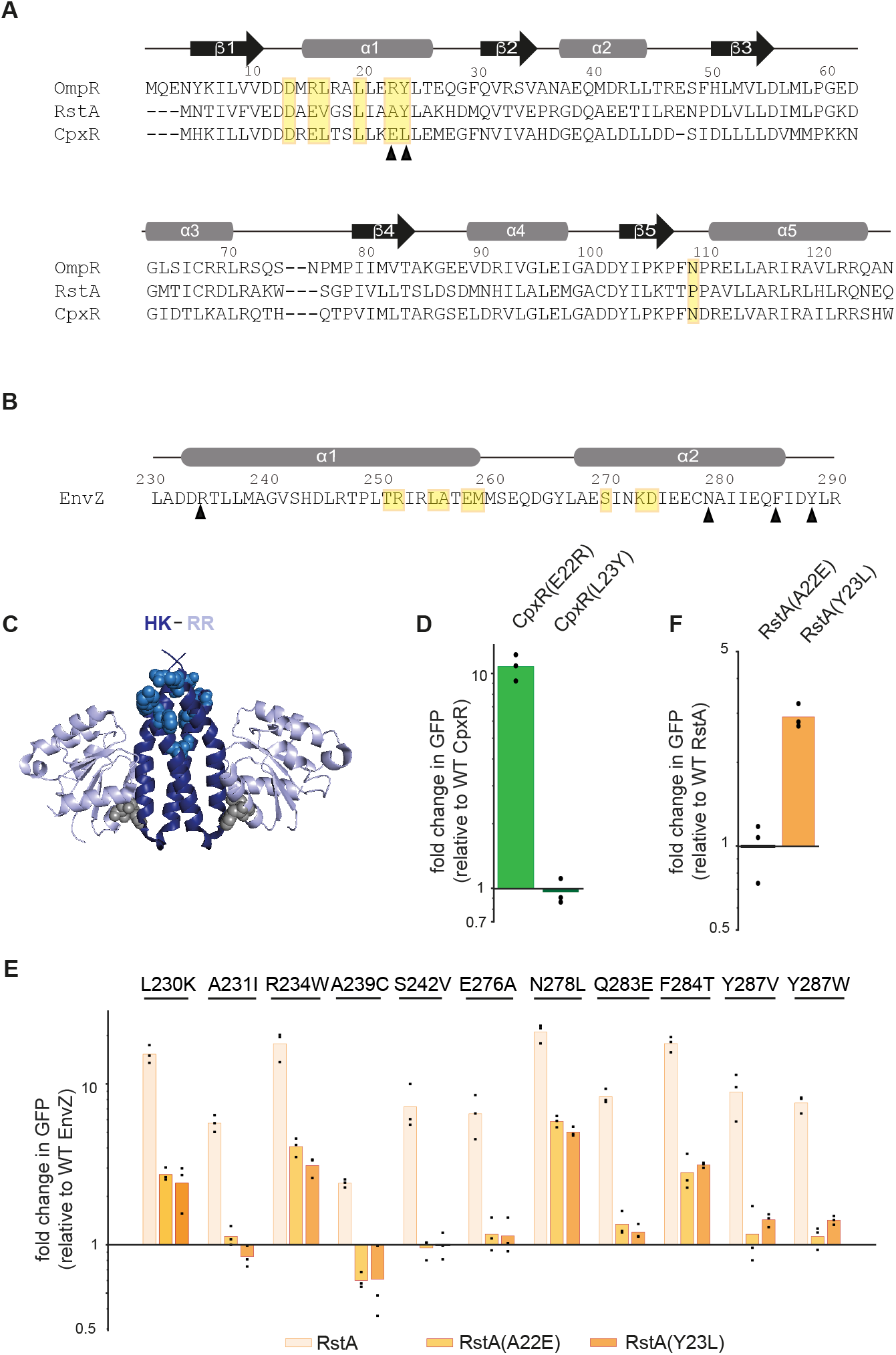
Additional characterization of RR substitutions. (A) Alignment of OmpR, RstA, and CpxR with coevolving residues highlighted in yellow. Arrowheads mark the two coevolving positions that differ significantly between CpxR and both OmpR and RstA. (B) EnvZ DHp domain sequence. Coevolving residues are highlighted in yellow. Arrowheads mark positions on EnvZ where secondary interface substitutions lead to cross-talk to RstA and that were tested for epistasis with RR substitutions. (C) Model of HK-RR complex with HK in deep blue and RR in light blue. Grey spheres on RR indicate coevolving residues of RstA and CpxR that were substituted and sky blue spheres on HK indicate distal residues of EnvZ that were substituted and tested against the variant RRs. (D) Fold changes in fluorescence of CpxR(E22R) and CpxR(L23Y) with wild-type EnvZ, relative to wild-type CpxR with wild-type EnvZ. n=3 biological replicates. (E) Fold changes in fluorescence relative to wild-type EnvZ for 7 additional distal single substitutions in EnvZ, against wild-type RstA, RstA(A22E), and RstA(Y23L). Data for 4 substitutions shown in Fig 4h are replicated here. n=3 biological replicates. (F) Fold changes in fluorescence of RstA(A22E) and RstA(Y23L) with wild-type EnvZ, relative to wild-type RstA with wild-type EnvZ. n=3 biological replicates. When compared to wild-type RstA, the substitutions tested in RstA do not lead to less phosphotransfer from wild-type EnvZ, suggesting that the decrease in activation by the EnvZ variants is not due to protein misfolding or lower expression.

**Fig. S8.**
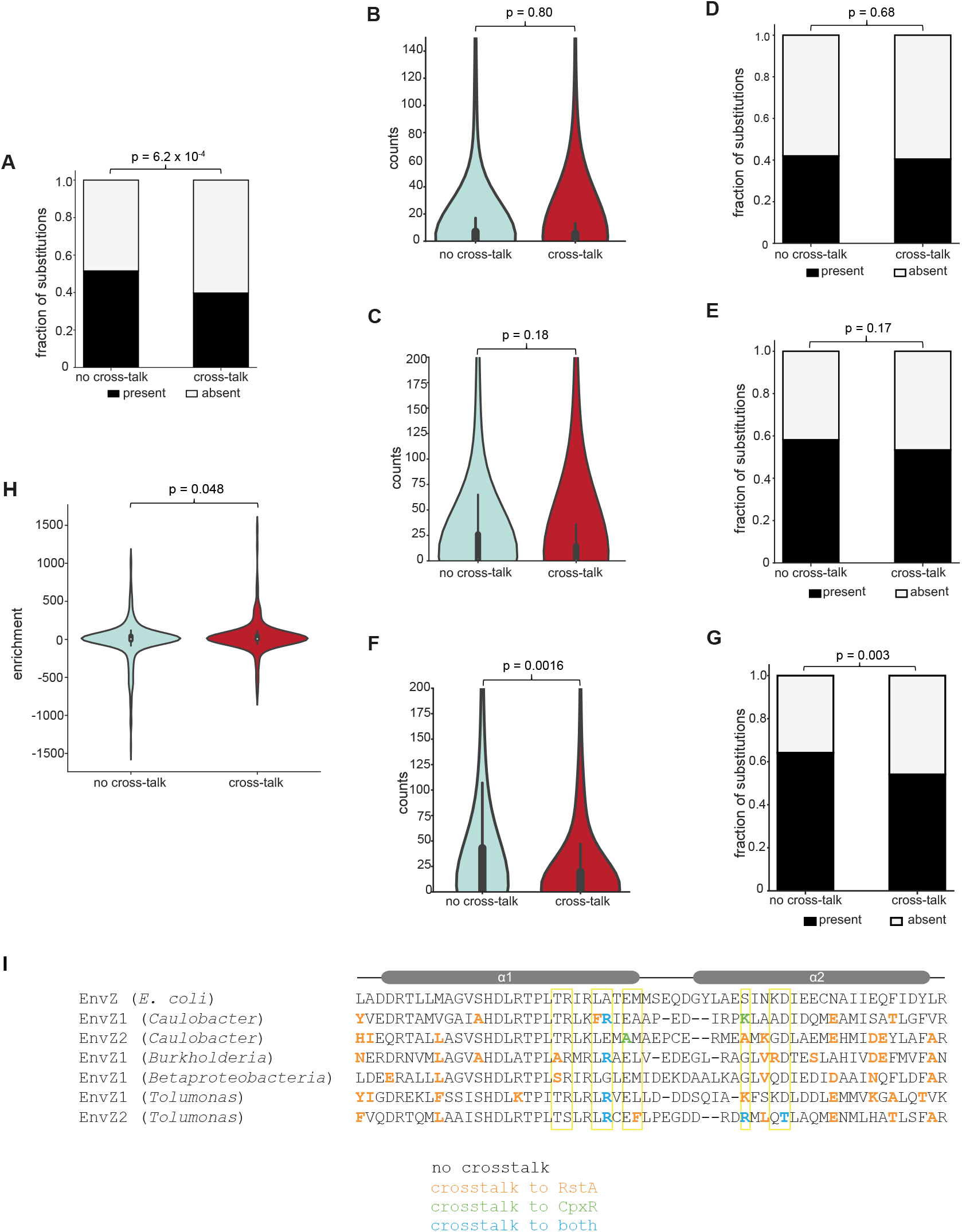
Additional bioinformatic analyses of histidine kinase ortholog sequences. (A) Fraction of substitutions observed to cross-talk or not (see Fig. 3) that are present or absent within the multiple-sequence alignment of 1019 EnvZ orthologs (p = 6.2 x 10^-4^, Fisher’s exact test). (B) Distributions of counts of EnvZ single substitutions that either do or do not cross-talk within a multiple sequence alignment of 594 YehU orthologs (p = 0.80, Kolmogorov-Smirnov test). (C) Same as (B) but for 1,088 BarA orthologs (p = 0.18, Kolmogorov-Smirnov test). (D) Same as (A) but for YehU (p = 0.67, Fisher’s exact test). (E) Same as (A) but for BarA (p = 0.17, Fisher’s exact test). (F) Same as (B) but for 1,067 PhoR orthologs (p = 0.0016, Kolmogorov-Smirnov test). (G) Same as (A) but for PhoR (p = 0.0030, Fisher’s exact test). (H) Violin plot showing distributions of enrichment of substitutions that either do or do not cross-talk in species that have lost RstBA and CpxAR, relative to species that retain RstBA and CpxAR (see Methods for enrichment calculation, p = 0.048, Kolmogorov-Smirnov test). (I) Alignment of EnvZ orthologs from species with duplications that have also lost RstBA and CpxAR, showing that these proteins have residues that cause cross-talk in the *E. coli* EnvZ background, as measured in our screen. Positions boxed in yellow are coevolving residues.

